# Spatial Profiling of Metals through Matrix-Assisted Laser Desorption Ionization Mass Spectrometry Imaging

**DOI:** 10.1101/2024.05.22.594980

**Authors:** Sylwia A. Stopka, Clément Bodineau, Gerard Baquer, Jia-Ren Lin, Md Amin Hossain, Michael S. Regan, Daniela Ruiz, Stecia-Marie Fletcher, Olivier Pourquié, Svetlana Lutsenko, Connor N. Payne, Jeffrey N. Agar, Ralph Mazitschek, Nathan J. McDannold, Peter K. Sorger, Sandro Santagata, Nathalie Y. R. Agar

## Abstract

The spatial-omic analysis of biomolecules such as nucleic acids, lipids, metabolites, and proteins is advancing the study of biological systems and processes in a physio-pathological context. Here, we describe an innovative matrix-assisted laser desorption ionization mass spectrometry imaging (MALDI MSI) method to detect metals within biological tissues using instrumentation that is widely available in research and clinical laboratories. We characterize the spatial distribution of metals in diverse settings including mouse embryogenesis, genetic disorders leading to abnormal metal accumulation, and preclinical testing for improved platinum-based chemotherapy delivery through focused ultrasound across the blood-brain barrier. Spatial metal profiling will advance research studies and the clinical analysis of metal-related diseases, enabling more precise use of metal-based therapies and advances in diverse scientific fields beyond biomedicine.

**One-Sentence Summary:** Spatial metallomic profiling maps native metals or those coordinated to xenobiotics, antibodies, and biomolecules in tissues.

## Introduction

Metallomics is an emerging field that focuses on the comprehensive study of metals in biological systems ^1^. Like other ‘-omics’ disciplines such as genomics, metabolomics, transcriptomics, and proteomics, the field of metallomics takes a systematic approach to investigate the roles of metals in normal cellular processes and their contributions to human diseases. The significance of the field is underscored by the fact that 30% of proteins in humans coordinate metal ions, including metalloenzymes, zinc finger proteins involved in DNA and RNA binding, copper-binding proteins, and iron-sulfur cluster proteins crucial for electron transfer reactions ^2-4^. In addition, imbalances in metal levels play a major role in numerous human diseases such as diseases resulting from toxic environmental exposures (e.g., lead, arsenic), neurodegenerative disorders like Alzheimer’s disease (AD) and Parkinson’s disease (PD) characterized by the accumulation of multiple metals in the brain, and genetic disorders like hemochromatosis and Wilson’s disease, which involve the abnormal accumulation of iron and copper in multiple organs ^5-7^.

Recent advances in spatial imaging methods have revolutionized the analysis of molecules within their native tissue context, providing valuable insights into the spatial organization of biomolecular processes. These technologies have expanded the dimensions of ‘-omics’ fields, leading to progress in spatial genomics, spatial metabolomics, spatial transcriptomics, and spatial proteomics as well as multi-modal approaches that combine these methods to yield datasets that integrate molecular information. One notable spatial profiling technology is matrix-assisted laser desorption/ionization mass spectrometry imaging (MALDI MSI) which enables the generation of quantitative maps of the distribution of small metabolites, lipids, proteins, and xenobiotics ^8-10^. MALDI MSI can be used for imaging the levels of drugs and their metabolites, as well as the molecular responses they induce in tissues from animal models and from patients enrolled in clinical trials ^11,12^. There is an increasing need to measure the spatial distribution of metals such as iron, copper, and calcium or metal-based drugs such as platinum-based chemotherapy agents (e.g., cisplatin, carboplatin, and oxaliplatin) as well as the gadolinium containing agents used in medical imaging by magnetic resonance imaging (MRI). Technologies such as atomic absorption spectrometry (AAS), inductively coupled plasma mass spectrometry (ICP-MS) ^13^, and secondary ion mass spectrometry (SIMS) ^14-16^ have been used to provide bulk measurements of metal levels, however, these methods lack detailed spatial information or provide poor spectral resolution.

Researchers have demonstrated the feasibility of detecting metals through MALDI MSI once they form chelate complexes with highly ionization-efficient derivatization reagents, such as dithiocarbamate (DDTC) ^17^. These DDTC-metal complexes enable the measurement of platinum using liquid chromatography-mass spectrometry (LC-MS) ^18,19^ and even the spatial mapping of its distribution ^20^. However, the formation of dimers and trimers by these complexes split the signal into multiple adducts, significantly limiting sensitivity and accurate quantification. We hypothesized that free metals, which have the advantage of already being ions in most biological settings, could be detected directly using MALDI-MSI. The widespread use of MALDI instruments in both scientific and clinical settings, along with their routine application for molecular tissue imaging, makes them an attractive option for research studies and diagnostics involving the direct spatial analysis of metals, metal containing compounds, and biomolecules.

## Exploring laser power and collision energy: two critical parameters to detect metals

To develop a method for the direct detection and imaging of metals (Fig. 1A) hypothesized that two key parameters must be carefully considered. First, the laser power (the energy of a UV pulsed laser in the MALDI ion source) would require optimization to achieve the desired level of gas-phase ion production, which requires releasing metals from their coordinated complexes in tissues and in some instances, forming detectable metal oxides (MO) ^21^. Second, the collision cell energy must be adjusted to fragment some ionized complexes, releasing intact free metals (M), allowing for their precise quantification and localization in tissues.

**Fig. 1.**
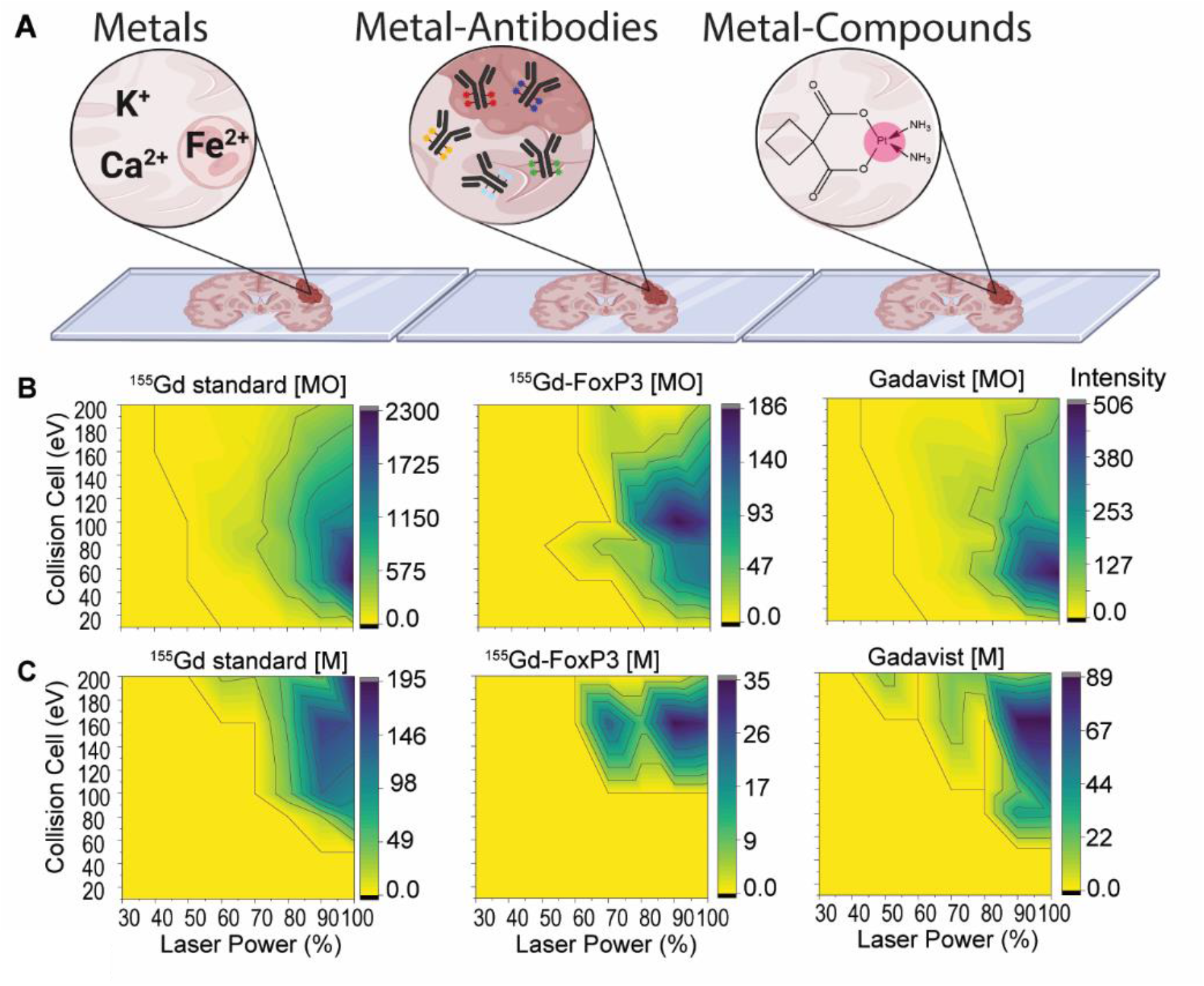
Workflow schematic for detecting metal and metal chelators using MALDI MSI is shown. **A**. For protein detection, frozen or formalin-fixed paraffin-embedded sections are stained with a panel of antibodies conjugated with isotopically pure metal chelator tags. For compounds containing metals such as proteins and free metals, no further sample preparation is needed, while for drugs and MRI imaging contrast dyes containing metals, the animals are first dosed with the compounds and sacrificed. The tissue of interest is then harvested and sectioned for analysis. At this point, both workflows converge and follow the same protocol. The tissues are sprayed with a MALDI matrix and imaged using a mass spectrometer with the instrument parameters tuned for metal detection. For the antibody workflow, within the same tissue, all the pure metal chelator tags can be detected, and their intensities spatially mapped over the tissue in the same run. For the drug and contrast dye dosed studies, the intensities of the metal chelators are also spatially mapped from the tissue. **B**. Heat maps illustrate the effect of collision cell energy and laser power on signal intensities of metal oxides and free metal ions.

To develop this approach, we first characterized the relationship between the UV laser energy (parameter 1) and the amount of gadolinium, a lanthanide metal, released from three different gadolinium-containing standards: a free ^155^Gd solution, a ^155^Gd-conjugated FoxP3 antibody, and Gadavist™, a gadolinium containing metal compound used as an MRI contrast agent. At lower levels of laser power, we observed baseline levels for free ^155^Gd. As we increased the laser power above settings typically used for organic molecules, we began to detect increasing amounts of the metal oxide (MO) form of gadolinium (Fig. 1B). In this context, the laser energy may act similarly to the high voltage energy used for declustering during electrospray ionization (ESI) ^19,22^. Increasing the collision cell energy (parameter 2) resulted in a shift of gadolinium from the metal oxide to the free metal (M) form (Fig. 1C). However, at ultra high collision cell energy levels, the signal was lost. This experiment demonstrated that optimization of both laser power and collision cell energy is required to detect metal oxides and free metal for each of the three standards. Using this approach suggested that direct metal analysis is achievable by MALDI mass spectrometry, eliminating the need for complex sample preparation steps such as those required for derivatization to detectable metallocompounds ^20^.

We next explored whether adjusting laser power and collision cell energy could enable the simultaneous imaging of multiple metals. We applied 11 free lanthanides (Tb, Eu, Pr, Sm, Gd, La, Nd, Lu, Tm, Er, Ho) onto a MALDI target plate and analyzed them after applying a range of collision energy (SI Fig. 1). In all 11 cases, we observed that higher laser powers resulted in stronger ion signals. However, for several free metals, we noticed that the ion signals plateaued at higher laser powers. In addition, while we found that the presence of MALDI matrix was not essential to detect free metals, it significantly increased the signal in many cases (by up to 250-fold). To explore the effect of varying collision energy on ion generation, we directly infused a platinum-chelated carboplatin complex into the mass spectrometer using ESI, ensuring no interference from the UV laser, and we monitored the levels of precursor carboplatin and platinum ions (SI Fig. 2). At lower collision energies, we detected a higher signal for the intact carboplatin complex. However, as the collision energy increased, the signal of free Pt metal substantially increased, while the ion intensity of the carboplatin-Pt complex decreased. This confirmed that the carboplatin complex bonds were dissociating, leading to the release of free Pt ions.

## Elemental mapping from preclinical and clinical biological tissue sections

After identifying instrument parameters that permit the measurement of multiple purified metals (SI table 1 and 2), we next focused on developing MALDI MSI to spatially profile metals within whole tissue sections. We acquired scans at varying rates, ranging from 1.5 min/mm^2^ at a100 µm pixel size to 153.8 min/mm^2^ at 5 µm (SI Table 3). This required further optimizing the relationship between laser power and collision energy, which we performed using 18 commercially available lanthanide-tagged antibodies. These antibodies were mixed with the MALDI matrix and applied to a stainless-steel surface. Optimization involved varying the laser power or collision cell energy on two separate mass spectrometers, both equipped with MALDI MSI capabilities – a quadrupole time-of-flight (SI Fig. 3-6) and a 15 Tesla Fourier-transform ion cyclotron resonance mass spectrometer (FTICR MS). Using both instruments, we successfully detected each lanthanide metal (Me) in its free form and within different complexes, such as metal oxides ([Me+O], [Me+OH], [Me+O_2_], and [Me+O_2_H_2_]) (SI Fig. 5-6).

We then used MALDI MSI to image a panel of five lanthanide-tagged antibodies applied to a formalin-fixed, paraffin-embedded (FFPE) human tonsil section at a 5-µm spatial resolution (Fig. 2A-C, 2E). Inspection of the images revealed the presence of immune, stromal, and epithelial markers in the expected compartments of the tonsil. The expression patterns of a subset of these markers were further confirmed using conventional multiplexed immunofluorescence imaging ^23^ performed on a serial section (Fig. 2D, 2F). Additionally, our MALDI-MSI data from the tonsil identified other metals such as Fe, Ca, Ni, and Cu, demonstrating the potential of combined multiplexed analysis of tagged antibodies and native metals (SI Fig. 7 and 8).

**Fig. 2.**
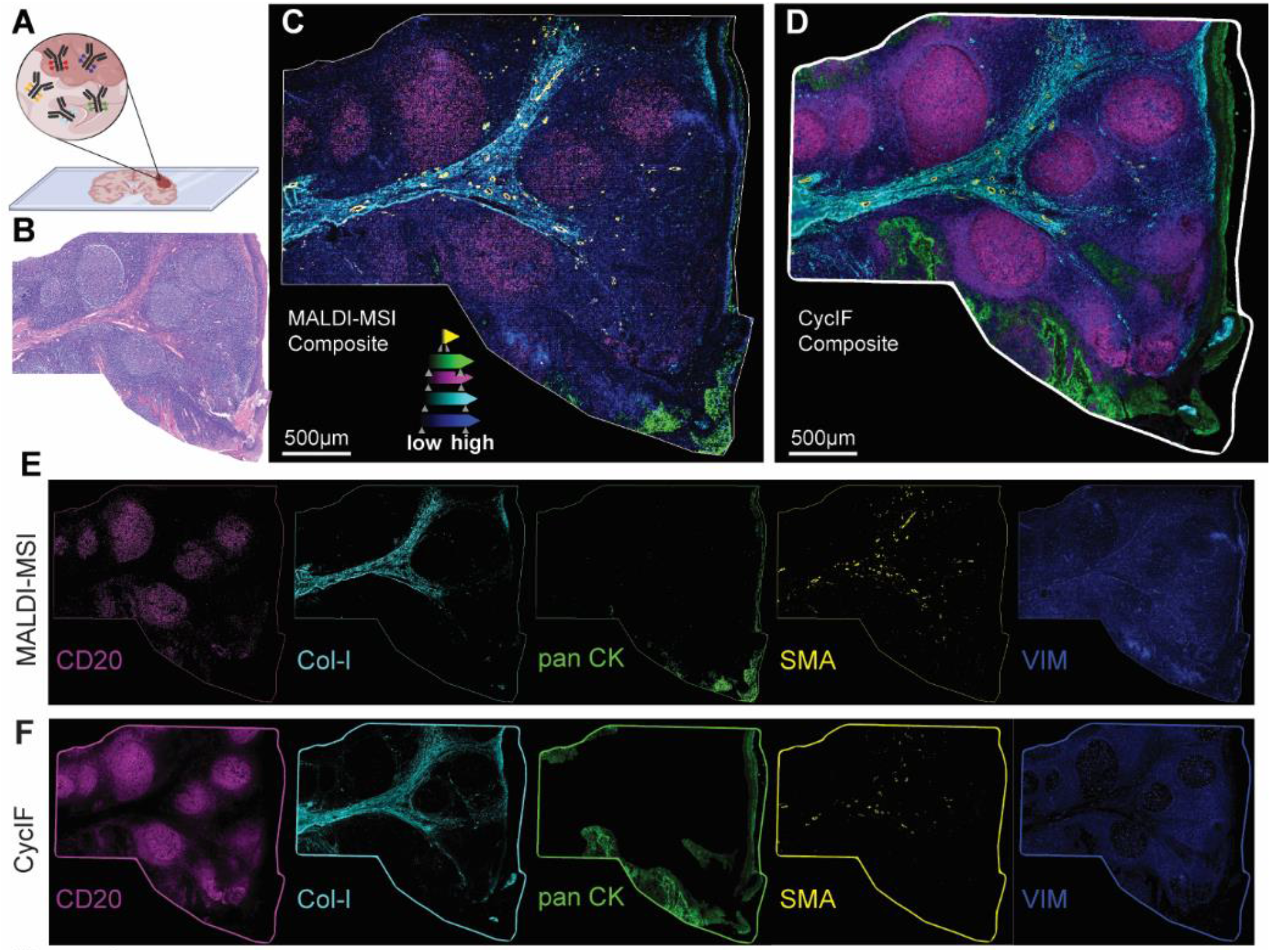
MALDI imaging of metal-labeled antibodies from human tonsil and validated by immunofluorescence. **A**. lanthanide-labeled antibodies. A brief schematic of lanthanide-tagged antibody metal analysis was provided. **B**. human tonsil H and E stain was used for the MALDI MSI. The tonsil was cryosectioned at 10 µm, fixed with paraformaldehyde, and incubated with the following metal-chelated antibodies: Smooth-muscle Actin 141Pr, Vimentin ^143^Nd, Pankeratin ^148^Nd, CD20 ^162^Dy, and Collagen ^169^Tm. It was then sprayed with 1,5-DAN and analyzed at a spatial resolution of 5 µm with a timsTOF mass spectrometer. **C** composite ion image of the five lanthanides that represent the antibodies was obtained. **D** Additionally, a composite image of a serial section that underwent cycIF for marker validation from tissue was obtained. **E-F** Individual ion images of the marker from MALDI and CycIF were also obtained.

We next focused on imaging native metals intrinsic to tissues in three different settings relevant to biomedical research and clinical diagnostics – (i) a mouse model of a human genetic disorder of impaired metal metabolism, (ii) a whole section of a developing mouse embryo just before birth, and (iii) whole sections of diagnostic biopsies from patients with an inherited disorder leading to metal overload (Fig. 3). Substantial amounts of copper are known to accumulate in the livers of mice with homozygous inactivation of the gene encoding *Atp7b*, a copper-transporting P-type ATPase mutated in Wilson disease ^24^. MALDI MSI of FFPE liver sections (∼0.5 cm^2^ of tissue area per specimen imaged at 50 µm spatial resolution) revealed an approximately 10-fold increase in copper levels in *Atp7b*-/- livers compared to those of wild type and *Atp7b*+/- mice (Fig. 3B).

**Fig. 3.**
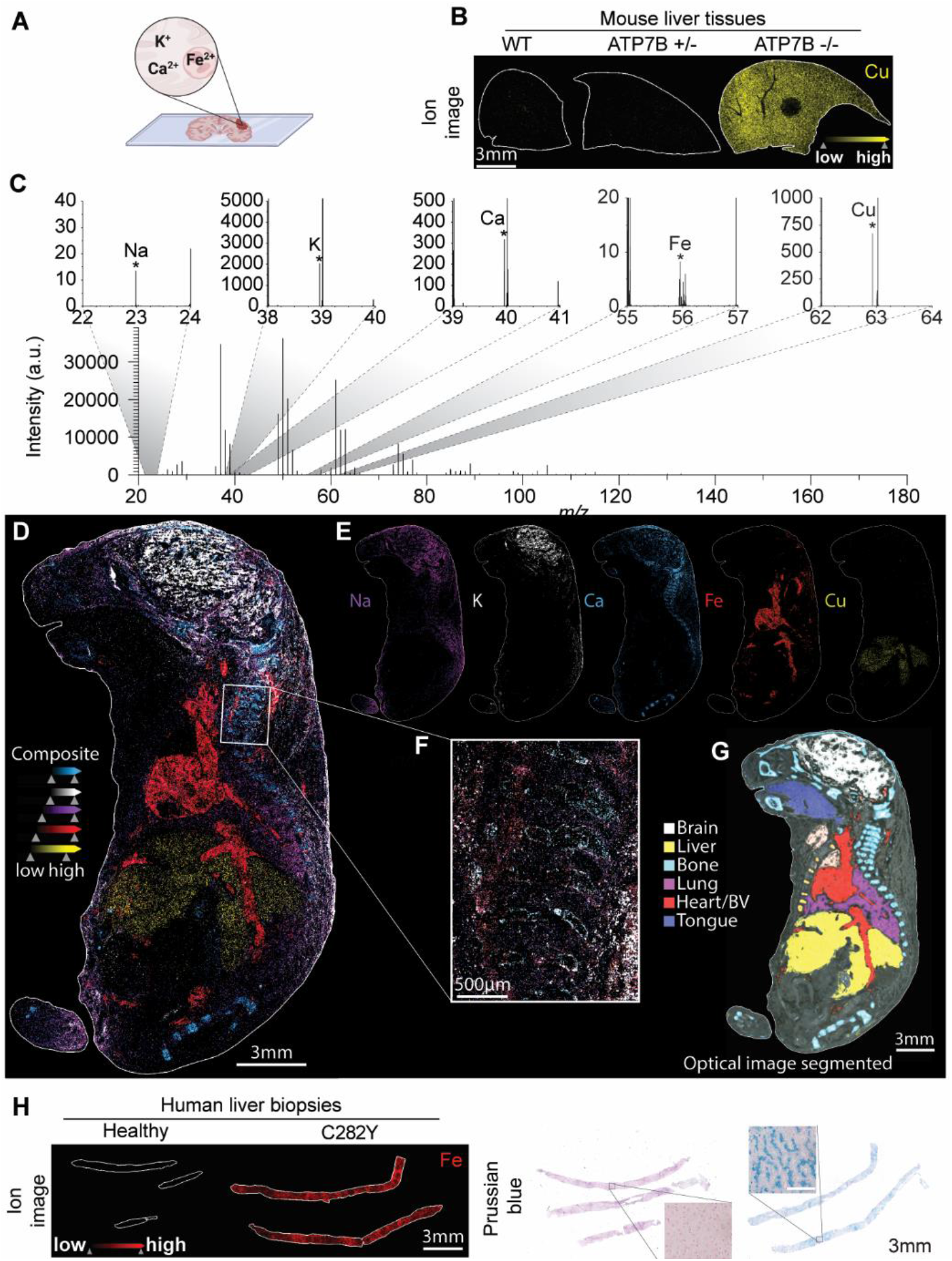
MALDI MS imaging of copper for Wilson disease, native metals from whole mouse embryo, and iron imaging from hemochromatosis diseased tissue. **A** brief schematic of free metal analysis. **B**. Iron distribution of a Wilson disease mouse model, where there is a 10-fold increase in iron signal compared to the WT. **C**. The overall spectrum and zoom-in regions highlight the detection of the metals. **D**. MALDI MSI of a whole mouse embryo at 20 µm with an ion image of E. five metals and their distribution in the tissue. **F**. The zoom-in is of a pixel size of 5 µm imaged area. **G**. An optical image of the embryo illustrates the different anatomical regions. H. Iron imaging of a hemochromatosis and control human liver biopsy, indicating a 1452 fold increase from normal to disease increase of iron. Prussian blue was used to confirm hemochromatosis.

In the second setting, MALDI MSI of a section of a whole mouse embryo (embryonic day E18.5; ∼3cm^2^ of tissue imaged at 20 µm spatial resolution) demonstrated the feasibility of imaging large tissue sections. A full scan spectrum revealed at least five distinct metals (Na, K, Ca, Fe, Cu) (Fig. 3D-F and SI Fig. 9). Segmentation of the organs and anatomical features based on an optical image of a serial section allowed us to quantify the relative levels of these metals, demonstrating distinctive metal distribution within each compartment (Fig. 3G). The localization of the metals matched the physiological functions; for example, sodium and potassium were highest in the brain, calcium in the bones, iron in the vascular system, and copper in the liver.

We detected the copper, iron, and sodium isotopes (SI Fig. 9) with relative abundances matching their known natural abundances (Table SI 1). For further validation, we imaged heme b from a normal mouse brain using previously reported MALDI-MSI methods ^25^. The overlapping signal for the iron and heme ions supports the specificity of the metal imaging results (SI Fig. 10).

In the third scenario, we imaged liver core needle biopsies obtained from patients with hemochromatosis, an inherited genetic disorder leading to excessive iron absorption and accumulation in multiple organs, including the liver. Scans of biopsies at 50 µm resolution showed a 1452-fold increase in iron in a patient homozygous for the *HFE* C282Y mutation (Fig. 3H). This provides a clear demonstration of the effectiveness of our method in detecting free metals in diverse contexts using MALDI MSI.

## Quantification of carboplatin and Gadavist in focused ultrasound treated tissue

To explore the application of spatial profiling of metals by MALDI MSI in guiding therapeutic development, we studied the effect of focused ultrasound (FUS) in temporarily and non-invasively opening the blood-brain barrier (BBB) in a pre-clinical rodent model of glioma for targeted delivery of carboplatin, a platinum-based chemotherapeutic agent, ^26^. We orthotopically engrafted a glioblastoma cell line in each hemisphere of a rat. Gadavist, a contrast agent containing the metal gadolinium, was used to assess the extent of transient BBB disruption.

After administering Gadavist and carboplatin (see methods section for details), FUS was applied selectively to open the BBB in one hemisphere only (Fig. 4A-B). MALDI MSI of metals from the frozen sections of the brain at a resolution of 20 µm allowed us to detect and quantify both Pt from carboplatin and Gd from Gadavist, as well as multiple native metals. Analysis of the whole tissue average spectrum revealed that the unique natural isotopic patterns of Pt and Gd aligned well with their known natural abundance (Fig. 4C-D). To quantify the levels of these compounds, we developed a tissue microarray mimetic ^27^ containing carboplatin and Gadavist in which the compounds were diluted and spiked into a tissue homogenate to generate standard calibration curves (SI Fig. 11). This approach revealed that the concentration of carboplatin in the sonicated tumor was 52 µM compared to 27 µM in the non-sonicated tumor. Similarly, the levels of Gadavist were also increased by approximately 2-fold, from 35 µM in the sonicated tumor versus 18 µM in the non-sonicated tumor (Fig. 4E-G). Using a modified MALDI MSI protocol, we were also able to measure and map the distribution of metals and metabolites from the tricarboxylic acid (TCA) cycle, glycolysis, and purinergic pathways from the same tissue section (Fig. 4H-J).

**Fig. 4.**
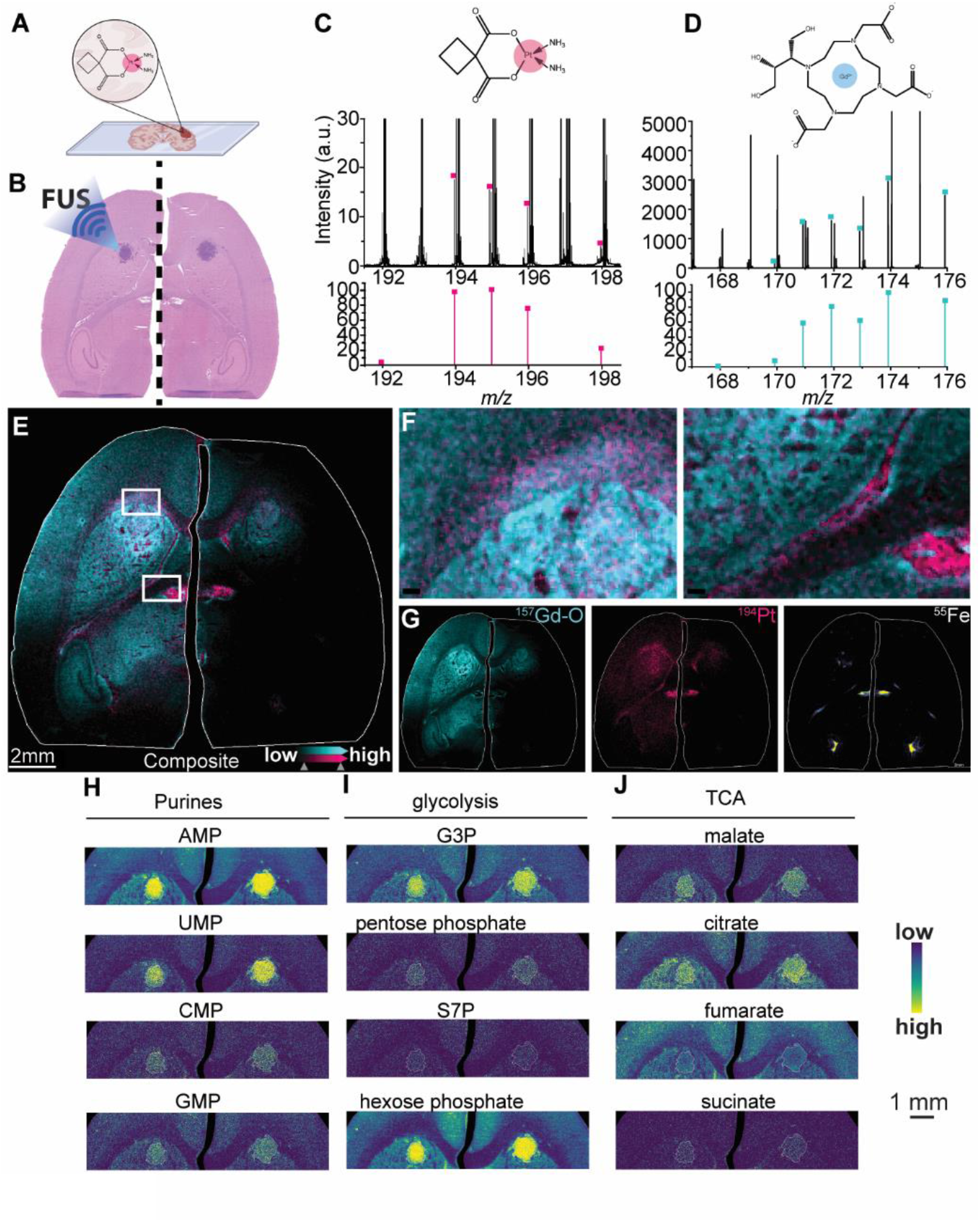
syngeneic glioblastoma tumor was orthotopically grafted into each hemisphere of a rat brain. The left hemisphere was treated with focused ultrasound, carboplatin, and gadavist to monitor blood-brain barrier opening, while the right hemisphere was not sonicated. **A**. A brief schematic analysis of metal compounds. **B**. A rat brain section was stained with H and E and used for MALDI MSI. **C-D** The averaged mass spectrum of the whole rat brain shows the unique isotope distribution of carboplatin and gadavist. **E-G** A composite image of both the drug and contrast dye was overlaid. The tissue was imaged for carboplatin and gadavist and then re-rastered with an MSI method tuned for small molecules, resulting in the ion images of several metabolites from the same tissue, including iron. **H**. ome small molecules detected from the same tissue and ion images for metabolites from the **I** purines, **J** glycolysis, and **K** TCA cycle.

In summary, we have developed an approach using MALDI MSI to image and quantify elements, revealing the distribution of metals, metal-containing drugs and MRI contrast agents, and metal/lanthanide-conjugated antibodies. The ability to rapidly and spatially profile metals presents many opportunities for preclinical and clinical applications and for advances in spatial metallomics. Beyond biomedical sciences and clinical use, our method holds potential for application in diverse fields such as environmental science, geology, material science, nanotechnology, and beyond.

## Methods

### Tissue collection

Liver biopsies from human formalin-fixed paraffin-embedded tissue samples were collected in accordance with the Institutional Review Board at Mass General Brigham (Partners Healthcare), Boston, MA, USA. Excess tissue discarded tissue protocol number: 2018P001627. Unfixed tonsil tissue was obtained and frozen by the Cooperative Human Tissue Network (CHTN). The Wilson disease mouse tissues were generously provided by Svetlana Lutsenko at Johns Hopkins. Specifically, ATP7B knockout mice (C57BL/6J-Atp7btm1Tcg/LtsnkJ), as previously described ^28^. All mouse handling procedures were conducted under the approval of the Johns Hopkins University Animal Care and Use Committee (JHU ACUC), protocol number MO15M27.

### Metal-tagged antibody tissue incubation

Flash-frozen or formalin-fixed, paraffin-embedded (FFPE) human tonsils were sectioned at 10 µm thickness on ITO-coated slides and fixed in PFA 4% at 4°C for 30 minutes. After permeabilization (twice 8 minutes in Triton-X100 0.2% in PBS at RT) and blocking (3% BSA in PBS for 45 minutes at RT), the sections were incubated with the antibodies overnight at 4°C in 0.5% BSA in PBS (Ki-67 (1:25, clone B56, metal168Er), collagen type I (polyclonal, 169Tm, 1:150), vimentin (1:50, clone D21H3, metal 143Nd)). Sections were washed twice for 8 minutes in Triton-X100 0.2% in PBS at RT, incubated with the Cell-ID Intercalator-Ir (1mM) for 30 minutes, and rinsed in HPLC grade water twice 5 minutes before desiccation.

### MALDI spectrometry Imaging of metals

The MSI data was acquired using a timsTOF flex and a 15 Tesla Bruker SolariX FT-ICR mass spectrometers (Bruker Daltonics, Billerica, MA) in positive ion mode to detect metals and negative ion mode for the detection of metabolites. On the timsTOF, the laser power was kept at 90% and the collision cell at 160 eV as this captured 22 of 35 metals, including Na, Mg, K, Ca, V, Cr, Mn, Fe, Ni, Co, Cu, Zn, Ga, Rb, Sr, Ag, Cd, Cs, Ba, Ti. Pb, and U using an ICP-MS (Multi-element Solution 2A, SPEX CertiPrep) tune mixture in which each metal at 10 µg/mL concentration (SI **Fig 7**). This mixture is also used to calibrate the mass range for MSI acquisition; using Rb, Na, Fe, and Cs peaks, the method can be calibrated from *m/z* 84.912 to 132.905. A mouse embryo was sectioned to 10 µm thickness and imaged using the new MSI method with the laser power at 90%, and the collision cell energy at 160 eV with a pixel size of 20 µm and a zoom-in area acquired with a 5 µm pixel size (SI **Fig 9**). Here the overall spectrum is shown as well as zoom-in *m/z* regions that highlight the different metals detected from the mouse embryo. The corresponding ion images of Na, K, Ca, Fe, and Cu and the localization within the tissue region are shown. Using the optical image, the organs and anatomical features were segmented and could be used to identify the unique metal distribution within each component.

For the lanthanide-antibody tags and Pt/Gd complexes, the mass range spans from 140.8947 to 208.947; thus, the mass range was calibrated using the Agilent tune mix solution (Agilent Technologies, Santa Clara, CA). To test this MSI approach, a human tonsil in FFPE was sectioned and incubated with Smooth-muscle Actin (^141^Pr metal), Vimentin 9 ^143^Nd metal), Pankeratin ^148^Nd CD20 (^162^Dy metal), and Collagen type 1 (^169^Tm metal). In this case, all metals were detected and spatially mapped (SI **Fig 8**). The validate this MALDI approach, cyclic immunofluorescence (CyCIF) was performed on a serial section and provided similar results.

### Cyclic immunofluorescence (CyCIF)

CyCIF staining was as described previously ^23^. In brief, the 5-micron FFPE tonsil sections were dewaxed on BOND RX autostainer (Leica Biosystems, Buffalo Grove, IL, USA), and antigen retrieval were performed with Leica ER1 buffer (Cat# AR9961) for 20 minutes. After dewaxing and antigen retrieval, the slides were immediately bleached with 4.5% H2O2 with LED light to reduce auto-fluorescent background. The antibodies were incubated overnight at 4°C. After washing, the slides were imaged with RareCyte CyteFinder imager using 20x Objectives. The multiple-cycle images was achieved via repetitively incubation-imaging-bleaching, followed by image stitching using ASHLAR package ^29^. The antibodies used were as follows: Collagen I (Polyclonal, Novus Biologicals, NB600-408AF647), pan-cytokeratin (AE1/AE3, ThermoFisher #41-9003-82), alpha-SMA (1A4, R&D #IC1420S), CD20 (L26, ThermoFisher #53-0202-82), Vimentin (D21H3, Cell Signaling #9856S).

### Carboplatin and gadavist dosed animals

All animal experiments were approved by the Institutional Animal Care and Use Committee (IACUC) at Brigham and Women’s Hospital. All methods were carried out in accordance with relevant guidelines and regulations. All study methods are reported in accordance with the Animal Research: Reporting of *In Vivo* Experiments (ARRIVE) guidelines.

Experiments were performed in male F98 tumor-bearing Fischer rats (∼250 g, Charles River Laboratories). Animals were anesthetized with an intraperitoneal (IP) injection of ketamine (80 ml/kg) and xylazine (10 ml/kg). Fur on the top of the head was removed using clippers and a depilatory cream.

F98 tumor cells were implanted bilaterally into the striata. Wild-type F98 cells (passage number five, CRL-2397, American Type Culture Collection) were cultured in high glucose Dulbecco’s Modified Eagle Medium (DMEM, 1×) supplemented with 10% Fetal Bovine Serum (FBS) and 5% Penicillin Streptomycin (P/S) in a humidified incubator with 5% CO2 at 37°C. In anesthetized rats, the dorsal surface of the skull was sterilized with an iodine swab. A 1 cm linear skin incision was placed over the bregma, and a 1 mm burr hole was drilled into the skull approximately 1 mm superior to and 3 mm lateral to the bregma. A 4 μL cell suspension (4.5 ×10^6^ cells/ml) was injected into the caudate putamen 3.5 mm from the dura surface using a 26-gauge, 10 μL airtight syringe (Hamilton). The cell suspension was injected at a rate of 0.9 µL/min using a digital syringe pump (Nanomite, Harvard Apparatus). Following the injection, the needle was allowed to sit for 3 minutes, before being slowly withdrawn over 3 minutes.

Animal behavior was monitored daily after surgery and sutures were removed after 5 days.

Focused ultrasound was delivered on day 9 after tumor implantation, when the tumor volumes measured using magnetic resonance imaging was ∼10 mm^3^. Prior to focused ultrasound treatment, the tail vein was catheterized with a 24-gauge catheter (Surflo®, Terumo) for intravenous (IV) administration.

Immediately following focused ultrasound blood-brain barrier opening, carboplatin (Millipore Sigma) was injected intravenously at a dose of 50 mg/kg over a period of approximately 90 s. At either 1, 4 or 24 hours after carboplatin administration, animals were sacrificed via cervical dislocation while under deep anesthesia. Fresh brains were flash-frozen in liquid nitrogen, and stored at -80°C.

### Focused ultrasound blood-brain barrier opening

Focused ultrasound blood-brain barrier opening was performed using a magnetic resonance imaging (MRI)-guided, clinical device (ExAblate Neuro, InSightec) that has been adapted for preclinical experiments. The device consists of a 30 cm-diameter,1024-element, steerable, hemispherical transducer array, a cavitation monitoring system, and a water system for cooling, degassing, and circulation. The operating frequency of the device is 220 kHz.

A clinical MRI scanner (Signa Premiere 3T, GE) was used to visualize tumors (T2-weighted fast spin echo) and for treatment planning (T2*-weighted Susceptibility Weight Angiography). An in-house constructed, rectangular receive-only surface coil (dimensions: 5×6 cm) was integrated with the system for imaging during the brain experiments. Following ultrasound sonication, a T1-weighted fast spin echo sequence was acquired before and after administration of MRI contrast (Gadavist, Bayer HealthCare Pharmaceuticals, 0.125 mmol/kg) to assess blood-brain barrier opening.

40 overlapping targets, with a 3 mm center-to-center spacing, were sonicated in a volume that covered most of the striatum (including the tumor and a surrounding margin) and most of the hippocampus in the right hemisphere. The contralateral hemisphere was not sonicated. Baseline sonications (total duration: 35 s) were first delivered without microbubbles for comparison to treatment sonications and to confirm a lack of cavitation activity. Sonications were then repeated with microbubbles for the actual treatment (total duration: 180 s). Microbubbles (Definity®, Lantheus Medical Imaging) were administered at the start of the sonication as a bolus injection through the tail vein at the approved clinical dose for ultrasound imaging, 10 µl/kg (∼1.2×10^8^ bubbles/kg). To facilitate the injections, the agent was diluted 10:1 in saline and was followed by a 200 µl injection of saline. FUS was delivered to the 40 target locations in the striatum over four volumetric sonications with 10 targets. Each volumetric sonication consisted of 5 ms bursts applied sequentially to the 10 targets with a pulse repetition frequency of 1.1 Hz interleaved over 180 s. A delay of at least three minutes ensured that most of the microbubbles had cleared before the start of the subsequent sonication.

Acoustic powers were modified using a real-time feedback controller, based on the analysis of acoustic emissions from cavitating MBs^30-32^. The starting acoustic power for sonications was 0.16 W, and the power was not allowed to exceed 2.58 W, which corresponded to estimated peak negative pressures in the range 102-444 kPa in water. It has previously been reported that transmission through the rat skull is in the range of 76-86% at a comparable frequency of 268 kHz ^33^. Passive monitoring of cavitation signals was performed using two receivers/hydrophones. The first was integrated into the ExAblate device and was resonant at the subharmonic frequency (115 kHz). The second was an in-house fabricated, elliptical (5 × 3 cm), air-backed receiver, with a resonant frequency of 660 kHz. A proportional feedback controller was used to independently modulate the power at each target during the exposures based on the strength of the second and third harmonic emissions at 440 and 660 kHz. The acoustic power was ramped until these signals reached a harmonic goal of 10-12 dB above the noise floor. The controller was designed to maintain harmonic emissions within this band for the first 30 s of the sonication, after which the power was set to the average power at which the harmonic goal was achieved during that time. If either wideband, subharmonic, or ultraharmonic (550 kHz) emissions were detected at any point during the sonication, the power was decreased by 15% and was not allowed to exceed that value for the remainder of the sonication. Detailed methods on this method of feedback control and methods for acoustic emissions acquisition with this set up have previously been described ^26,31^.

### Quantitative metal MALDI MSI

For Pt and Gd metal quantification from carboplatin and gadavist, respectively, a tissue mimetic was constructed using homogenized control mouse brain tissue spiked with the two compounds. An eight-point nonzero calibration point was used for carboplatin were the homoygenized tissue were spiked at final concentrations of 50 μM, 10 μM, 5.0 μM, 2.0 μM, 1.0 μM, 0.8 μM, 0.2 μM, 0.1 μM, 0.0 μM, while a six nonzero calibration point for gadavist included 50 μM, 20 μM, 10 μM, 5.0 μM, 3.0 μM, 1.0 μM, and 0.0 μM (SI **Fig 11a-b**). The spiked tissue homogenates were then dispensed into a pre-casted 40% gelatin tissue microarray consisting of 1.5 mm core diameter wells, and the whole array was frozen until analysis. The quantitative tissue mimetic arrays were then cryo-sectioned at ten μm thickness and placed onto the same ITO slide containing the carboplatin and gadavist dosed tissue sections. After MSI acquisition, the data were processed and visualized using SCiLS lab software (version 2023a core, Bruker Daltonics, Billerica, MA) without normalization (SI **Fig 11c-d**). An average ion intensity of either Pt (*m/z* 194.963) or Gd (*m/z* 173.917) was collected from each calibration point for both carboplatin and gadavist, and a linear relationship between ion intensity and analyte concentration was plotted. Using the linear regression, the concentration of carboplatin and gadavist could be calculated in any region, *e*.*g*., sonicated or non-sonicated tumor within the rat brain. For each analyte, the correlation coefficient (R^2^), the limit of quantification (LOQ) with an S/N ratio of > 10, and the limit of detection (LOD) with an S/N ratio of >3 were calculated. For carboplatin the Pt quantitation was observed to be linear within 50-0.0 μM with an R^2^ = 0.9998, LOQ = 2.3 μM, and LOD = 0.70 μM. For Gadavist, the Gd quantitation was linear within 50-0.0 μM with an R^2^ = 0.9906, LOQ = 19.02 μM, and LOD = 5.71 μM. Additionally, the tissue can be reimaged for metabolite spatial mapping; for example, the rat brain tissue was reimaged, and metabolites such as glutamate and heme b could be mapped on the same tissue section (SI **Fig 12**).

## Acknowledgments

Acknowledgments follow the references and notes list but are not numbered. Start with text that acknowledges non-author contributions and then complete each of the sections below as separate paragraphs.

## Author contributions

Conceptualization: NYRA, SAS, CB, NJM

Methodology: NYRA, SAS, CB

Investigation: NYRA, SAS, CB, GB, JRL, MSR

Visualization: NYRA, SAS, CB, GB, JRL, MSR

Funding acquisition: NYRA

Project administration: NYRA

Supervision: NYRA, OP

Writing – original draft: NYRA, SAS

Writing – review & editing: NYRA, SAS, CB, GB, JRL, MSR, DR, OP, SS, SMF, NJM

## Data and materials availability

The data and materials associated with this study are available upon request. Inquiries regarding access to the data, datasets, code, or any other materials used in this research can be directed to Professor Nathalie Y.R. Agar at.

## Supplementary Materials

Materials and Methods

Figs. S1 to S12

Tables S1 to S3

