## Supplementary Information for "Spatial Profiling of Metals through Matrix-Assisted Laser Desorption Ionization Mass Spectrometry Imaging"

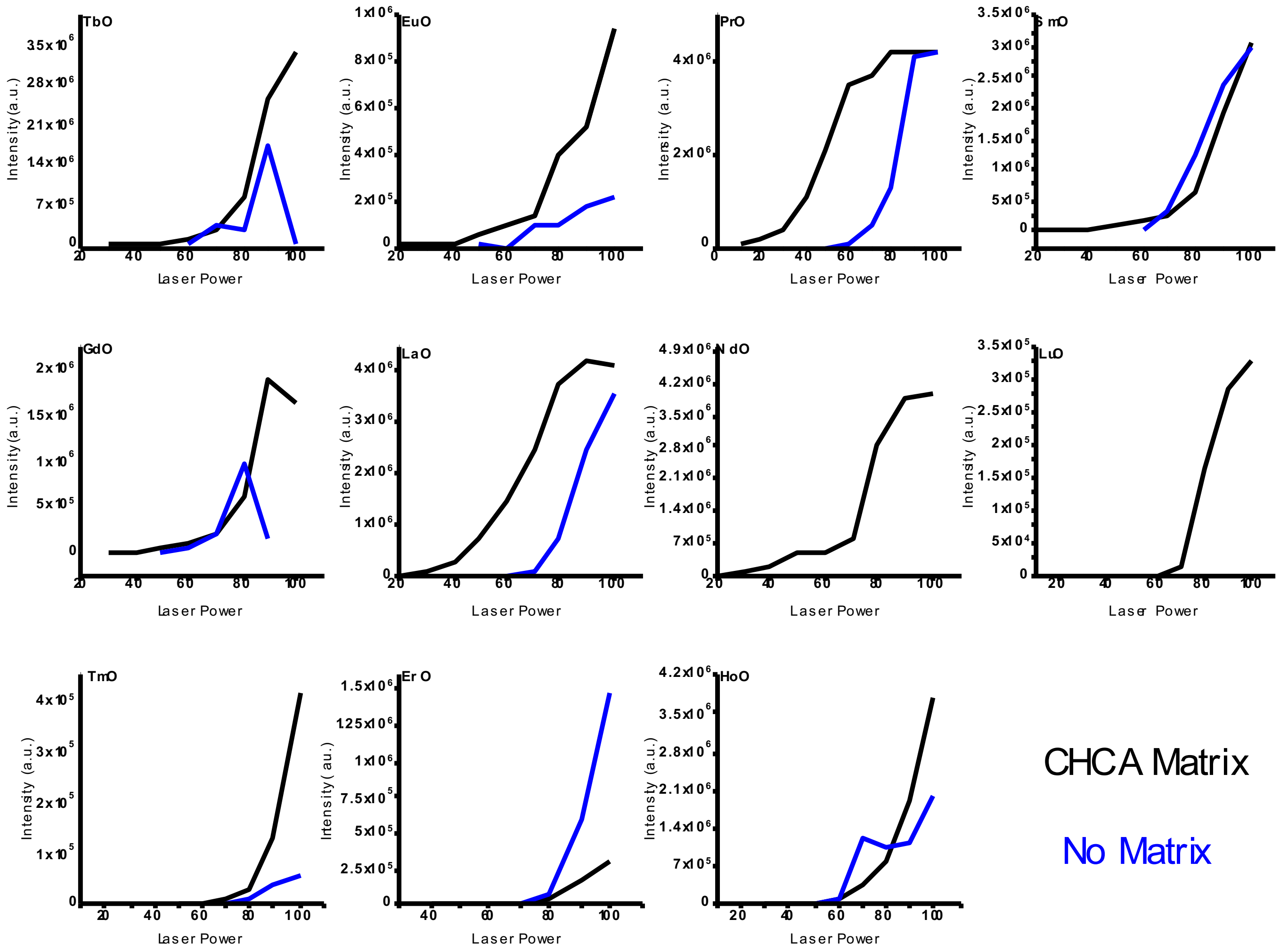


**SI Figure 1.** Free lanthanide metal detection with and without the assistance of a MALDI matrix. Using a timsTOF mass spectrometer, Tb, Eu, Pr, Sm, Gd, La, Nd, Lu, Tm, Er, and Ho were spotted onto a MALDI target plate with CHCA and without any matrix. The laser power was varied from 10 to 100% in which 10,000 laser shots were averaged.

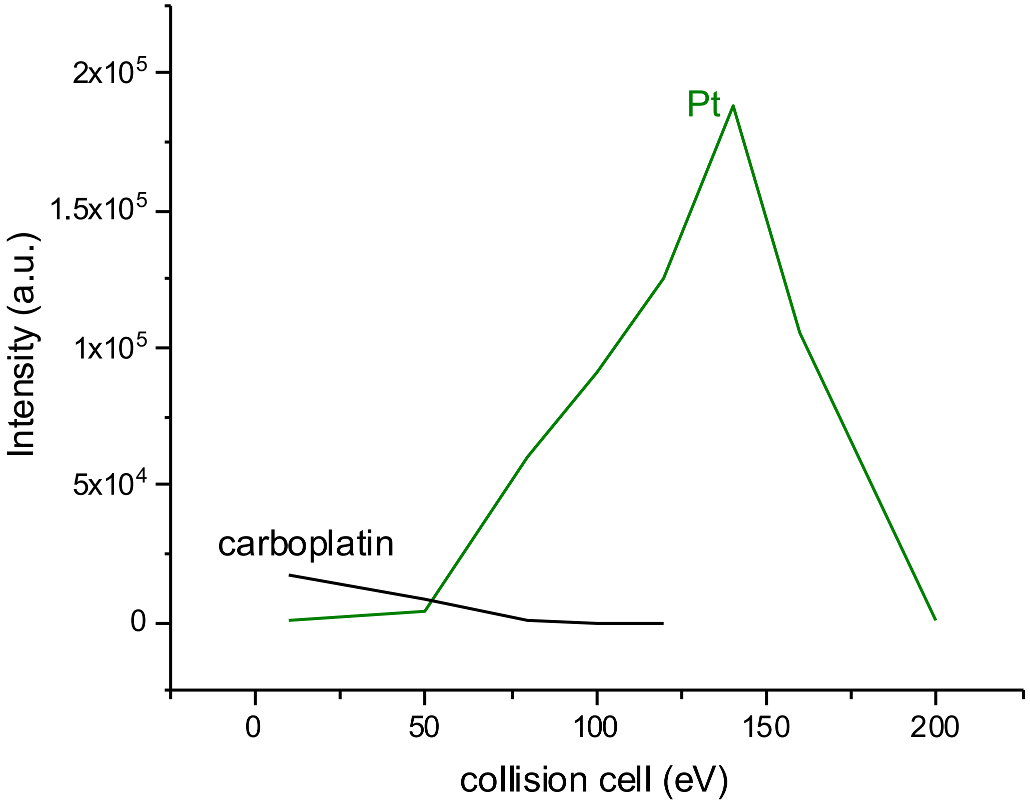


**SI Figure 2.** timsTOF mass spectrometry analysis of carboplatin as the laser power is set to a constant 90% power with varying collision cell (CC) energy. The carboplatin complex decreases in intensity as the CC energy is increased. Monitoring the Pt at *m/z* 194. 965 peak shows an increase of signal as the CC energy is increased.

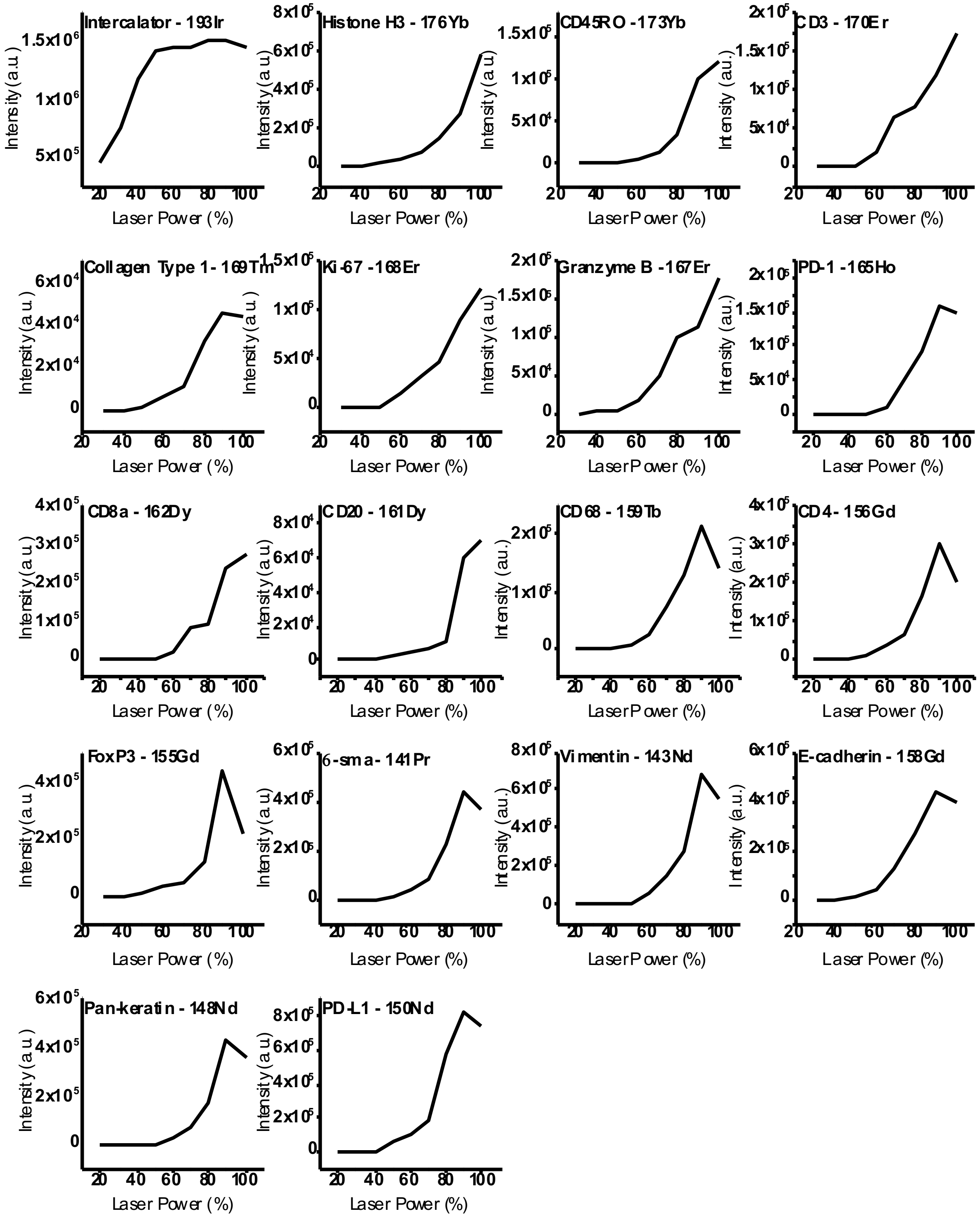


**SI Figure 3.** MALDI MS using a timsTOF mass spectrometer of 17 individual standard antibodies and a cationic nucleic acid intercalator conjugated with rare earth metals with varied laser powers. The collision cell energy was kept at a constant 160 eV while the laser was ramped from 20 to 100% power.

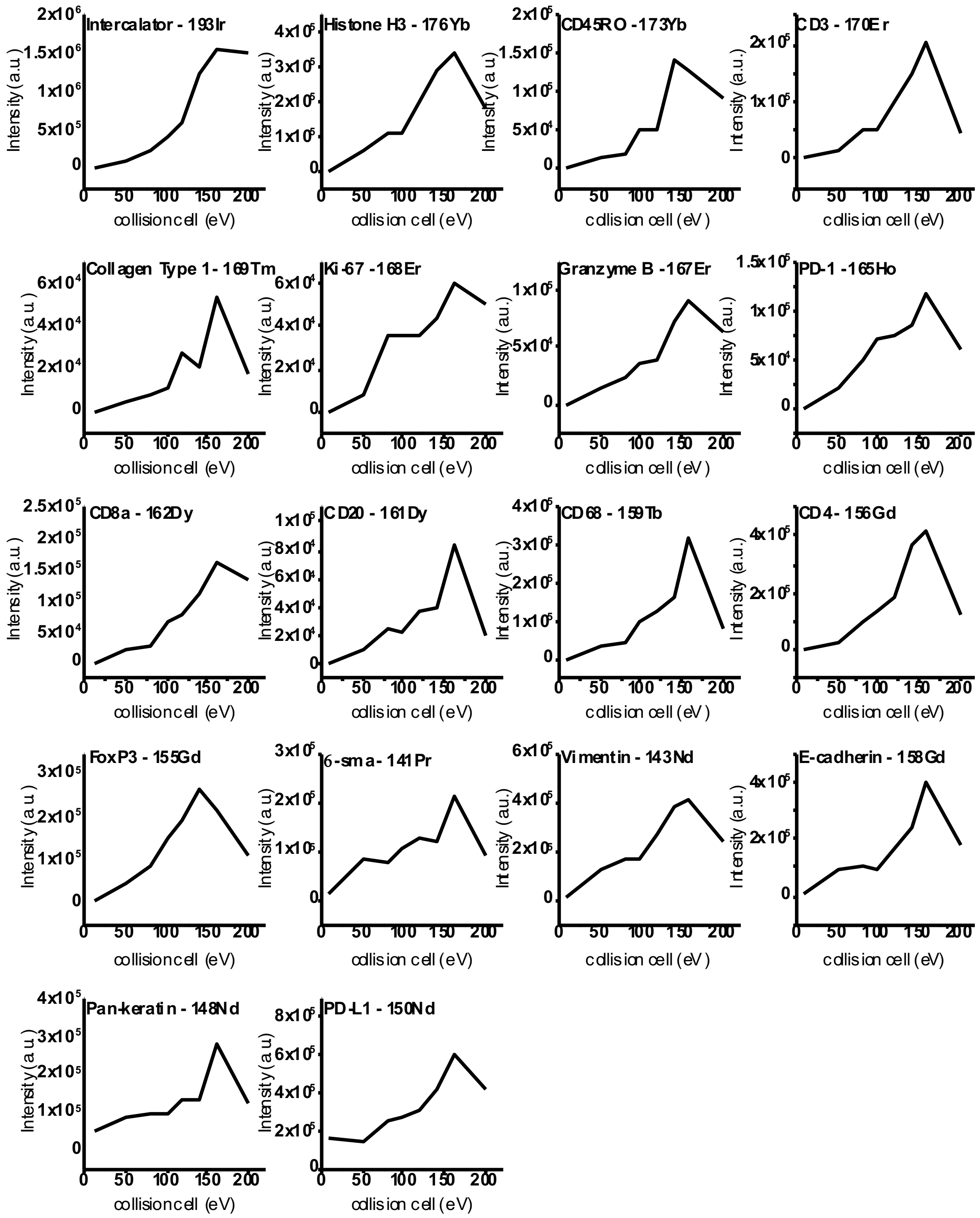


**SI Figure 4.** MALDI MS using a timsTOF mass spectrometer of 17 individual standard antibodies and a cationic nucleic acid intercalator conjugated with rare earth metals with varied collision cell energies. The laser power was kept at a constant 90% while the collision cell energy was ramped from 10- 200 eV.

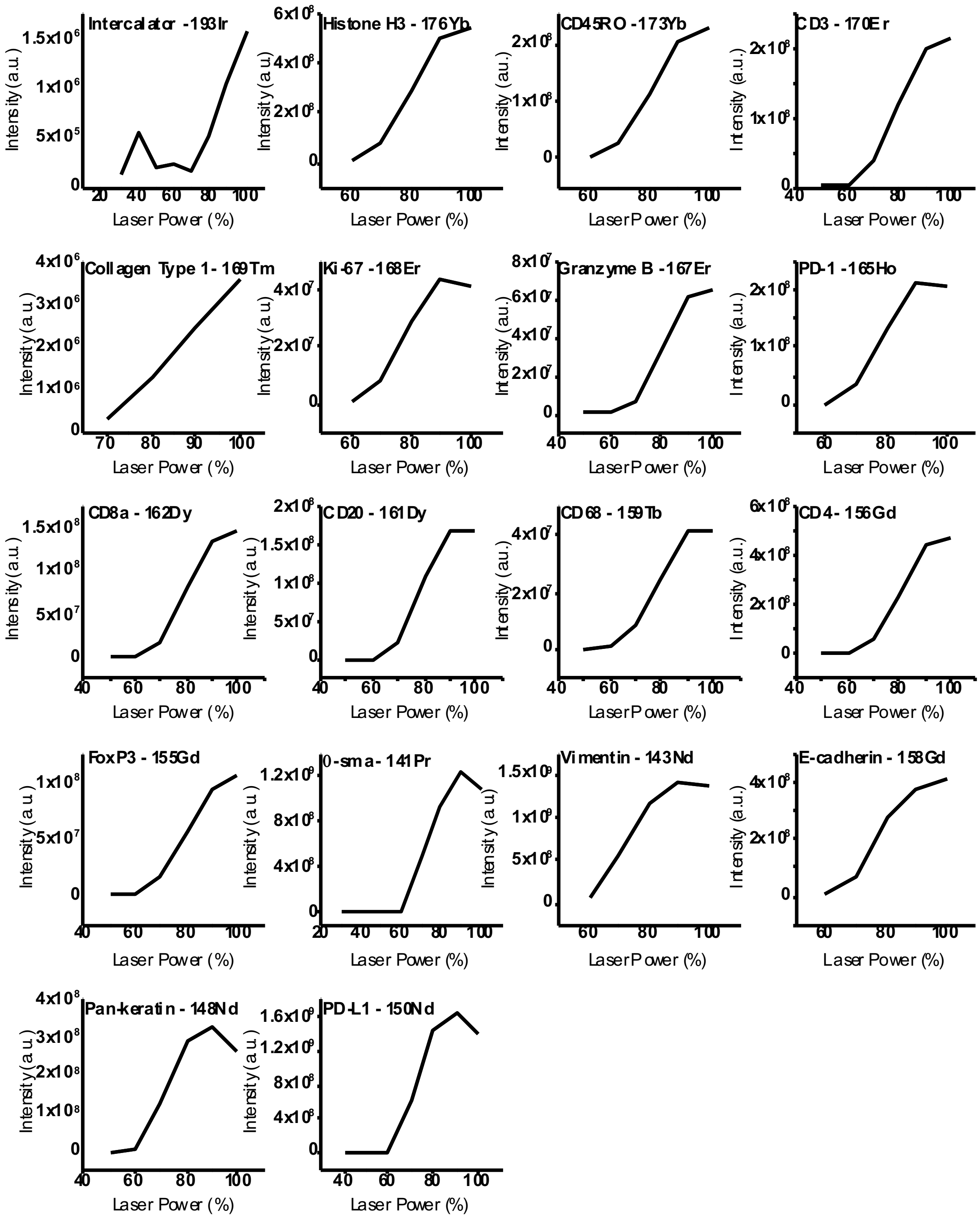


**SI Figure 5.** MALDI MS using a 15T FT-ICR mass spectrometer of 17 individual standard antibodies and a cationic nucleic acid intercalator conjugated with rare earth metals with varied laser powers. The collision cell was kept at a constant 20 V while the laser was ramped from 30 to 100% power.

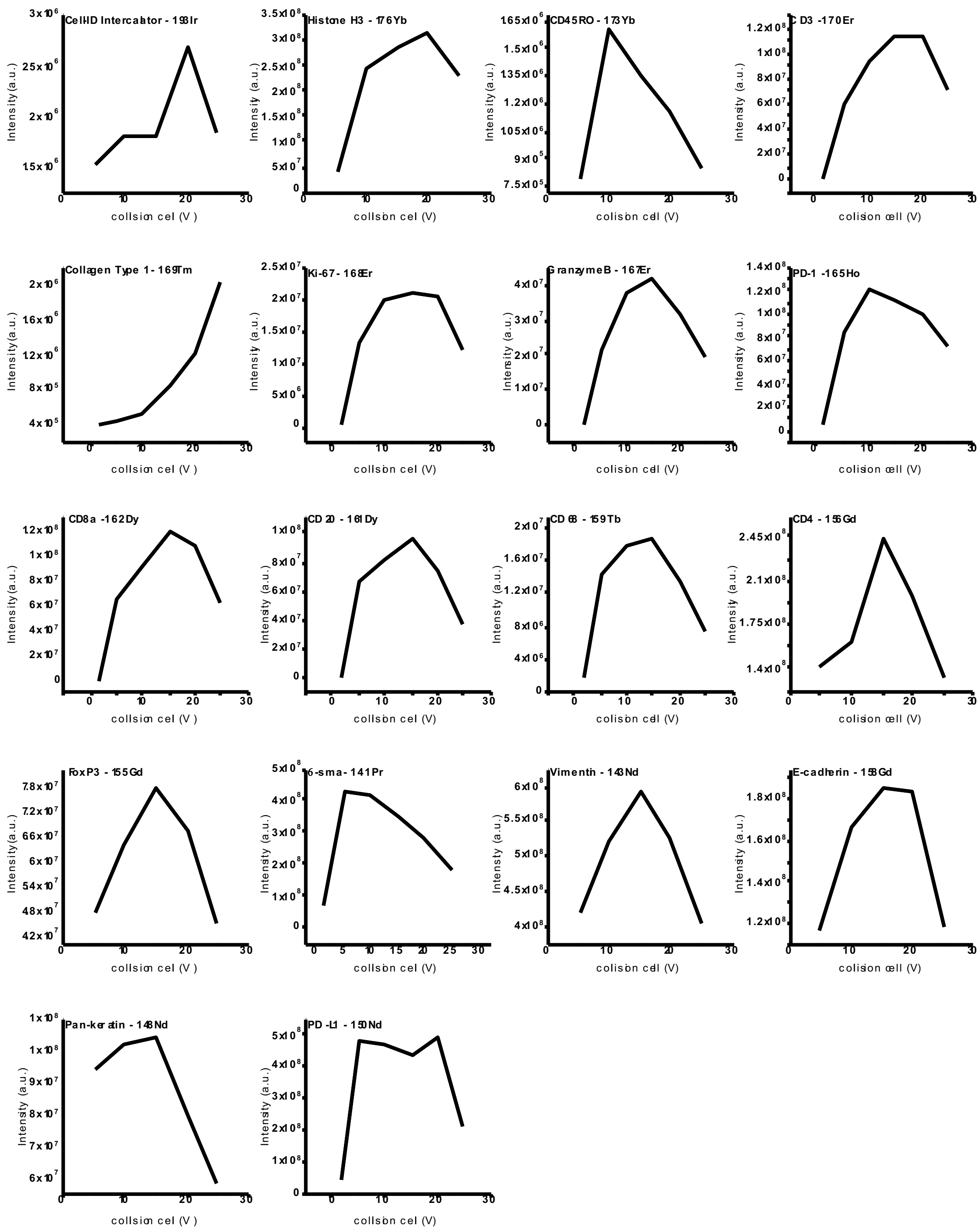


**SI Figure 6.** MALDI MS using a 15T FT-ICR mass spectrometer of 17 individual standard antibodies and a cationic nucleic acid intercalator conjugated with rare earth metals with varied collision cell energies. The laser power was kept at a constant 90% power while the collision cell energy was ramped from 1.5-25 V.

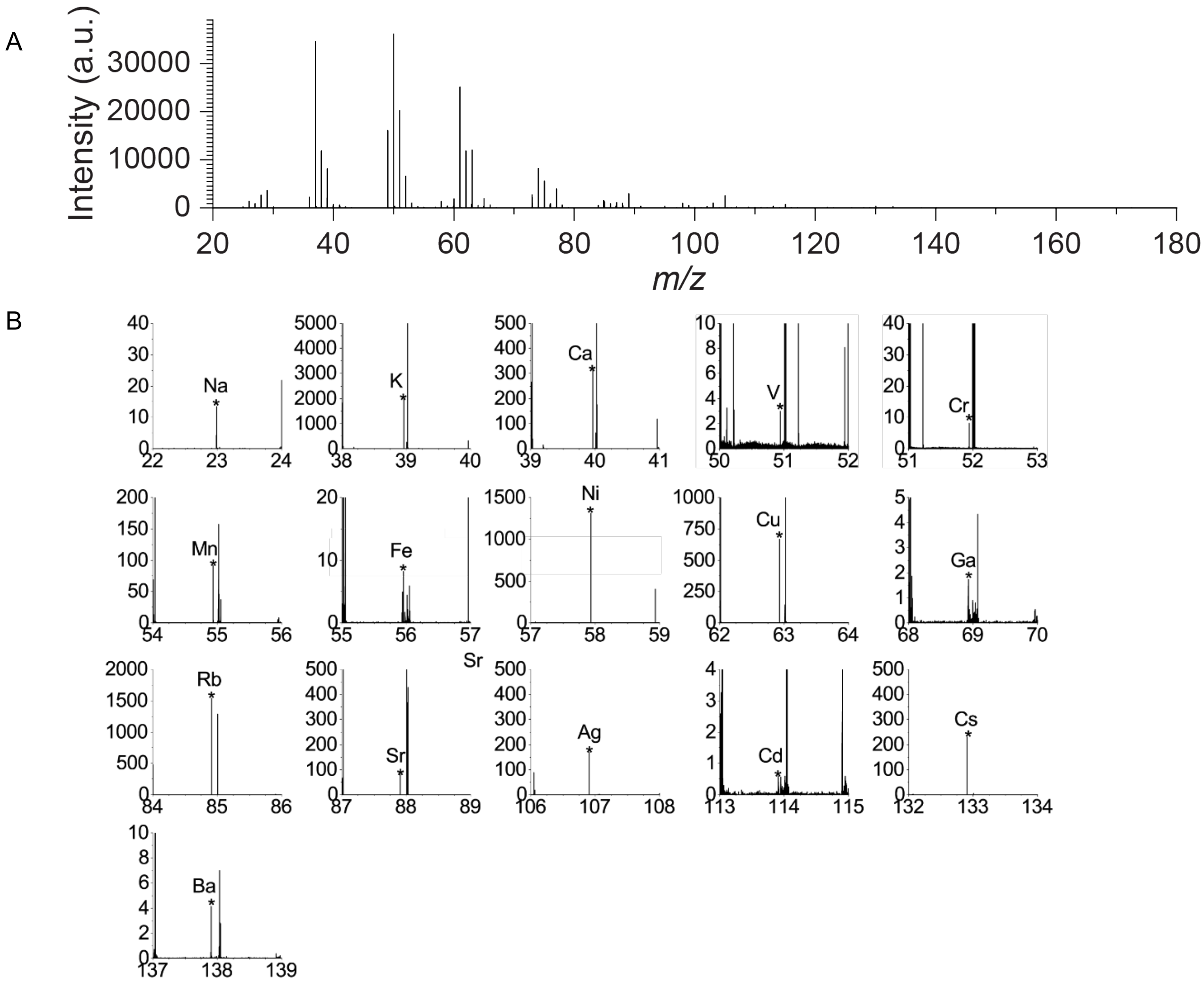


**Figure SI 7. (A)**MALDI MSI droplet analysis of a metal standard mixture in positive ion mode on a Q-ToF mass spectrometer. **(B)** Zoom-in regions of individual metals from the metal standard mixture.

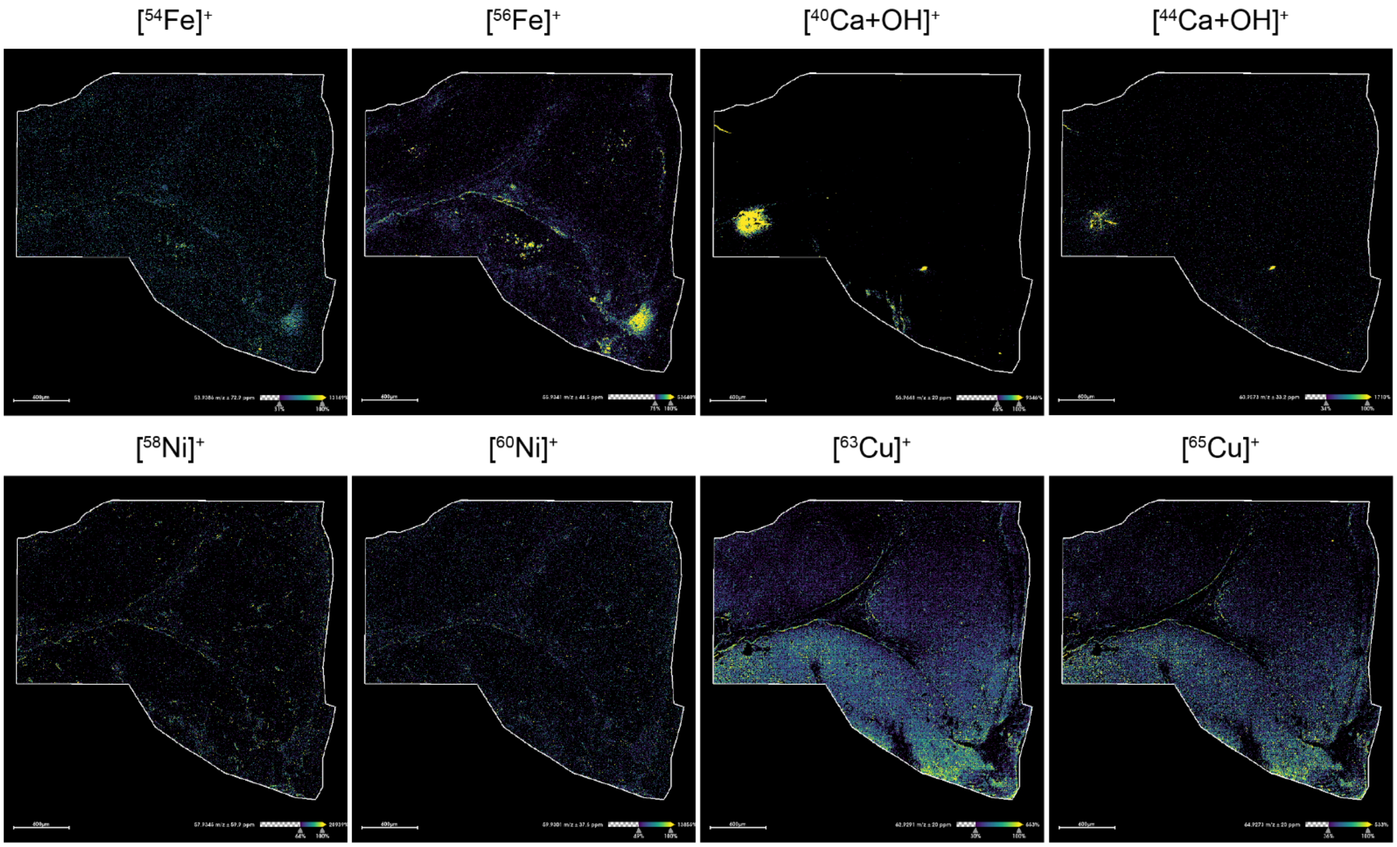


**Figure SI 8.** MALDI-MSI of elemental endogenous metals from a serial section of human tonsil from Fig 2 and collected on a Q-ToF at 5 µm spatial resolution.

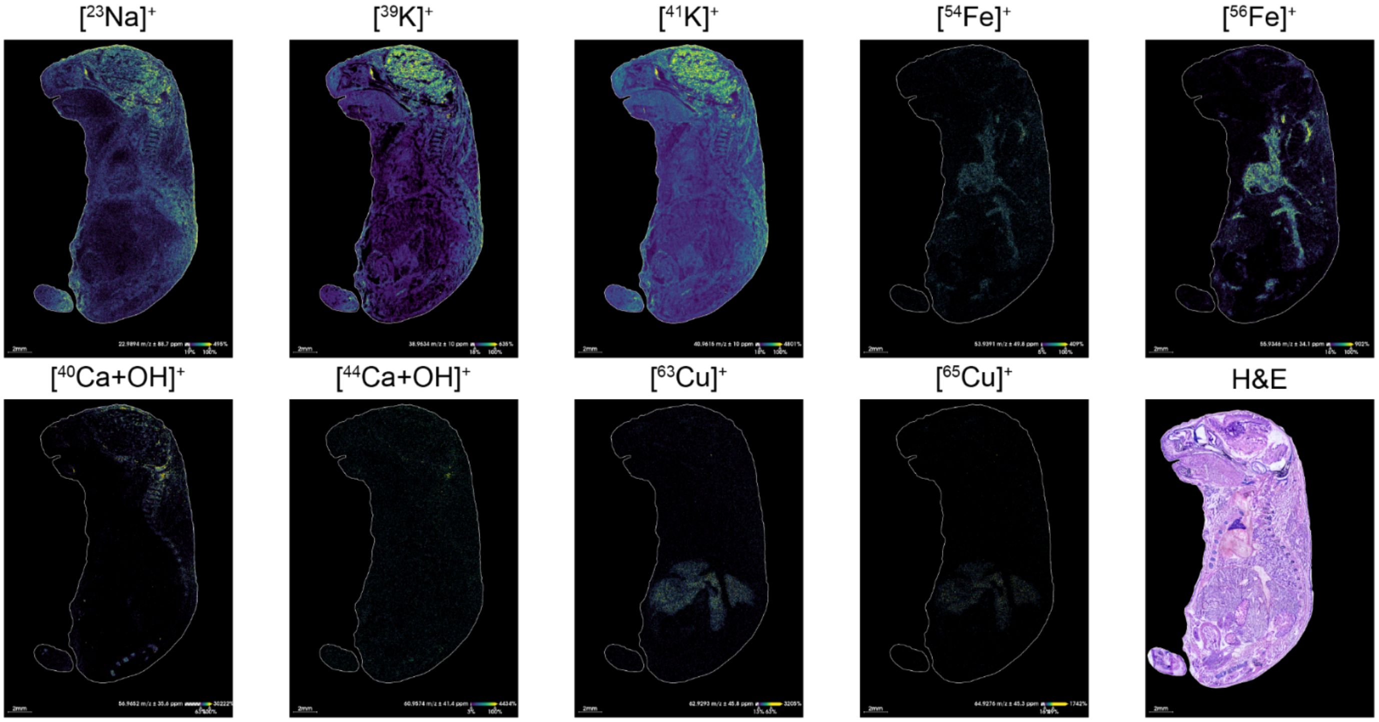


**SI Figure 9.** Additional ion images and H&E image of elemental endogenous metals from the whole mouse brain from Fig 3 and collected on a Q-ToF at 20 μm spatial resolution.

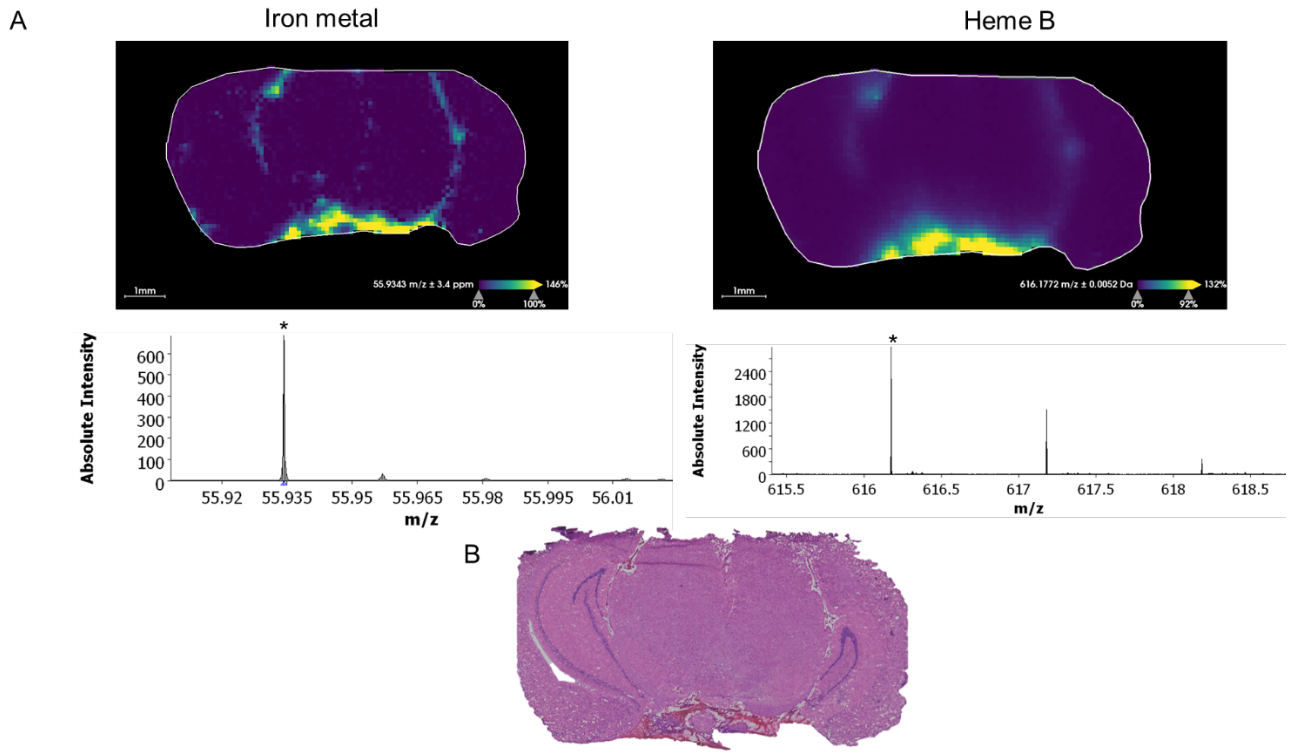


**SI Figure 10.** (A) MALDI MSI of coronal mouse brain tissue with 1,5-DAN matrix applied. Tissue was scanned for the distribution of iron (left), then rescanned with another method to see the distribution of heme b. Both ion images have a similar pattern. * on the spectra indicates the peak of respectively Fe at *m/z* 55.935 and heme B at *m/z* 616.178. (B) H&E staining of the section after MALDI MSI acquisition showing correlation between the ion image and the vasculature of the tissue.

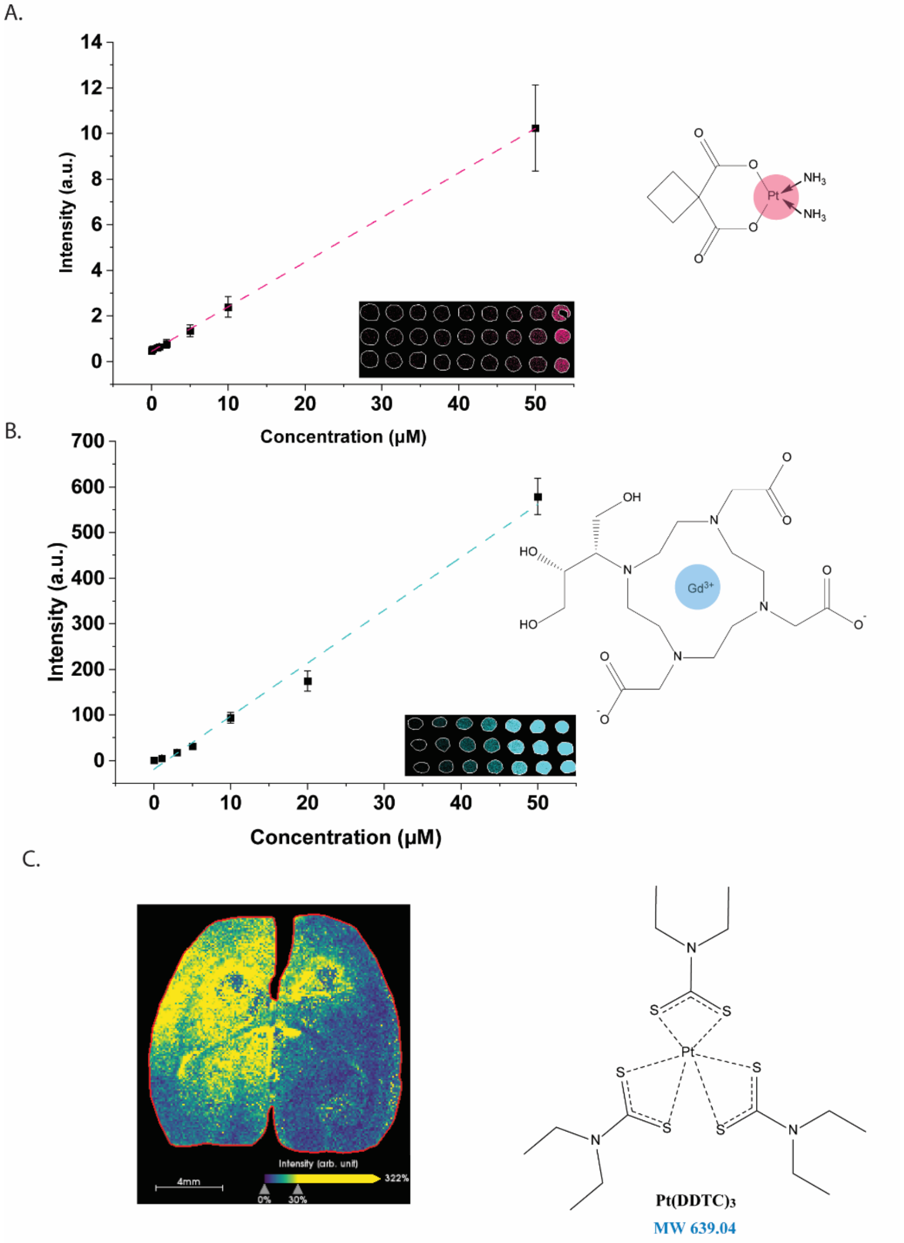


**SI Figure 11.** A rat brain was orthotopically grafted with a syngeneic glioblastoma tumor in each hemisphere, co-treated with carboplatin and gadavist to monitor blood-brain barrier opening following focused ultrasound treatment of the left hemisphere compared to the non-sonicated right hemisphere. Using a timsTOF mass spectrometer, quantitative MALDI MSI response of (A) carboplatin and (B) gadavist by monitoring Pt and Gd, respectively. (C) validation of Pt imaging of a serial tissue section that underwent on-tissue chemical derivatization using diethyldithiocarbamate (DDTC).

| Target | Clone | Metal | $[M+O]_{Theoretical}$ | $[M+O]_{Experimental}$ | PPM |
|---|---|---|---|---|---|
| Alpha-smooth muscle actin | 1A4 | $^{141}Pr$ | 156.9020 | 156.9021 | 0.36 |
| Vimentin | D21H3 | $^{143}Nd$ | 158.9042 | 158.9042 | 0.34 |
| Pan-keratin | C11 | $^{148}Nd$ | 163.9113 | 163.9113 | 0.21 |
| PD-L1 | SP142 | $^{150}Nd$ | 165.9153 | 165.9153 | 0.32 |
| FoxP3 | PCH101 | $_{155}Gd$ | 170.9170 | 170.9170 | 0.26 |
| CD4 | EPR6855 | $^{156}Gd$ | 171.9165 | 171.9165 | 0.25 |
| E-cadherin | 24E10 | $_{158}Gd$ | 173.9185 | 173.9185 | 0.36 |
| CD68 | KP1 | $_{159}Tb$ | 174.9197 | 174.9198 | 0.88 |
| CD20 | H1 | $^{161}Dy$ | 176.9213 | 176.9213 | 0.19 |
| CD8a | C8/144B | $^{162}Dy$ | 177.9212 | 177.9212 | 0.16 |
| PD-1 | EPR4877(2) | $^{165}Ho$ | 180.9247 | 180.9248 | 0.36 |
| Granzyme B | EPR20129-217 | $^{167}Er$ | 182.9264 | 182.9265 | 0.27 |
| Ki-67 | B56 | $^{168}Er$ | 183.9267 | 183.9268 | 0.09 |
| Collagen Type I | Polyclonal | $^{169}Tm$ | 184.9286 | 184.9286 | 0.03 |
| CD3 | Polyclonal, C-terminal | $^{170}Er$ | 185.9298 | 185.9298 | -0.03 |
| CD45RO | UCHL1 | $^{173}Yb$ | 188.9326 | 188.9327 | 0.68 |
| Histone H3 | D1H2 | $^{176}Yb$ | 191.9369 | 191.9370 | 0.20 |

**SI table 1**. List of Maxpar antibodies that were detected using a 15 T FT-ICR MS using MALDI

| | | ESI | | MALDI | |
|---|---|---|---|---|---|
| Metal | $[M]_{Theoretical}$ | $[M]_{Experimental}$ | PPM | $[M]_{Experimental}$ | PPM |
| Li | 7.0160 | - | - | - | - |
| Be | 9.0121 | - | - | - | - |
| Na | 22.9897 | 22.9898 | 4.35 | 22.9896 | -4.35 |
| Mg | 23.9850 | 23.9851 | 4.17 | 23.9849 | -4.17 |
| Al | 26.9815 | 26.9816 | 3.71 | 26.9816 | 3.71 |
| K | 38.9637 | 38.9639 | 5.13 | 38.9636 | -2.57 |
| Ca | 39.9625 | 39.9628 | 7.51 | 39.9625 | 0.00 |
| V | 50.9439 | 50.9439 | 0.00 | 50.9435 | -7.85 |
| Cr | 51.9405 | 51.9405 | 0.00 | 51.9405 | 0.00 |
| Mn | 54.9380 | 54.9380 | 0.00 | 54.9380 | 0.00 |
| Fe | 55.9349 | 55.9350 | 1.79 | 55.9349 | 0.00 |
| Ni 58 | 57.9353 | 57.9351 | -3.45 | 57.9353 | 0.00 |
| Co | 58.9331 | 58.9330 | -1.70 | 58.9332 | 1.70 |
| Ni 60 | 59.9307 | 59.9307 | 0.00 | 59.9308 | 1.67 |
| Cu 63 | 62.9295 | 62.9295 | 0.00 | 62.9296 | 1.59 |
| Zn | 63.9291 | 63.9289 | -3.13 | 63.9293 | 3.13 |
| Cu 65 | 64.9277 | 64.9277 | 0.00 | 64.9277 | 0.00 |
| Ga | 68.9255 | 68.9252 | -4.35 | 68.9255 | 0.00 |
| As | 74.9215 | 74.9214 | -1.33 | 74.9214 | -1.33 |
| Se | 79.9165 | 79.9165 | 0.00 | 79.9164 | -1.25 |
| Rb | 84.9117 | 84.9118 | 1.18 | 84.9117 | 0.00 |
| Sr | 87.9056 | 87.9056 | 0.00 | 87.9056 | 0.00 |
| Mo 96 | 95.9046 | 95.9047 | 1.04 | 95.9045 | -1.04 |
| Mo 97 | 96.9060 | 96.9055 | -5.16 | 96.9053 | -7.22 |
| Mo 98 | 97.9054 | 97.9054 | 0.00 | 97.9052 | -2.04 |
| Mo 99 | 99.9074 | 99.9078 | 4.00 | 99.9078 | 4.00 |
| Ag | 106.9050 | 106.9051 | 0.94 | 106.9052 | 1.87 |
| Cd | 113.9033 | 113.9031 | -1.76 | 113.9031 | -1.76 |
| Cs | 132.9054 | 132.9050 | -3.01 | 132.9055 | 0.75 |
| Ba | 137.9052 | 137.9050 | -1.45 | 137.9053 | 0.73 |
| Hg 198 | 197.9667 | 197.9651 | -8.08 | 197.9663 | -2.02 |
| Hg 199 | 198.9682 | 198.9686 | 2.01 | 198.9697 | 7.54 |
| Hg 200 | 199.9683 | 199.9678 | -2.50 | 199.9687 | 2.00 |
| Hg 201 | 200.9703 | 200.9698 | -2.49 | 200.9710 | 3.48 |
| Hg 202 | 201.9706 | 201.9701 | -2.48 | 201.9710 | 1.98 |
| Hg 204 | 203.9734 | 203.9726 | -3.92 | 203.9738 | 1.96 |
| Tl | 204.9744 | 204.9738 | -2.93 | 204.9745 | 0.49 |
| Pb | 207.9766 | 207.9761 | -2.40 | 207.9772 | 2.88 |
| U | 238.0507 | 238.0497 | -4.20 | 238.0511 | 1.68 |

**SI table 2**. List of metals that were detected using a Q ToF using MALDI and ESI

| min/mm$^2$ | pixel size (µm) |
|---|---|
| 0.4 | 100 |
| 1.5 | 50 |
| 10.4 | 20 |
| 37.7 | 10 |
| 153.8 | 5 |

**SI Table 3**. Experimental scanning rates of the timsTOF (Bruker) MSI instrument using the metal detecting method where the laser frequency was set to 10,000 Hz and the pixel size was varied from 5 to 100 µm.
